# Integrative Omics Analysis Reveals the Immunomodulatory Effects of the Parasitic Dinoflagellate *Hematodinium* on Crustacean Immunity

**DOI:** 10.1101/2021.11.17.468922

**Authors:** Meng Li, Qian Huang, Xiaoyang Lv, Hamish J. Small, Caiwen Li

## Abstract

Parasitic dinoflagellates in genus *Hematodinium* have caused substantial economic losses to multiple commercially valuable marine crustaceans around the world. In the present study, comprehensive omics approaches (miRNA transcriptomics, iTRAQ-based proteomics) were applied to investigate the host-parasite interaction between hemocytes from *Portunus trituberculatus* and *Hematodinium perezi*. The parasitic dinoflagellate remodeled the miRNome and proteome of hemocytes from challenged hosts, modulated the host immune response at both post-transcriptional and translational levels and caused post-transcriptional regulation to the host immune response. Multiple important cellular and humoral immune-related pathways (ex. Apoptosis, Endocytosis, ECM-receptor interaction, proPO activation pathway, Toll-like signaling pathway, Jak-STAT signaling pathway) were significantly affected by *Hematodinium* parasites. Through modulation of the host miRNome, the host immune responses of nodulation, proPO activation and antimicrobial peptides were significantly suppressed. Cellular homeostasis was imbalanced via post-transcriptional dysregulation of the phagosome, peroxisome, and lysosome pathways. Cellular structure and communication was seriously impacted by post-transcriptional downregulation of ECM-receptor interaction and focal adhesion pathways.

**Author summary:** The parasitic dinoflagellate *Hematodinium* infects many economically important marine crustaceans. Recent efforts to better understand the life cycle and biology of the parasite have improved our understanding of the disease ecology. However, studies on the host-parasite interaction, especially how *Hematodinium* parasites evade the host immune response are lacking. To address this shortfall, we used miRNA transcriptomics and iTRAQ-based proteomic approaches to explore the immune responses of *Portunus trituberculatus* when challenged with *Hematodinium perezi*. Striking changes in the miRNome and proteome of hemocytes were observed, and the parasite exhibited multifaceted immunomodulatory effects and potential immune-evasion mechanisms in this crustacean host.

## Introduction

Dinoflagellates are a phylum of unicellular eukaryotes, functioning as the dominant primary producers in the oceanic and coastal ecosystem besides diatoms [1]. Taxonomically, dinoflagellates belong to the infrakingdom Alevolata that also comprise the phyla Apicomplexa and Ciliophora, and form a monophyletic group close to apicomplexan parasites [2, 3]. Dinoflagellates consist of photosynthetic and non-photosynthetic species, with approximately half of the known species being mixotrophic, heterotrophic or parasitic [4, 5]. Both ecto- and endoparasitic dinoflagellates are common in marine environments, infecting hosts ranging from ciliates, radiolarians and planktonic dinoflagellates to crustaceans, cnidarians and many others [6]. A recent exploration of marine eukaryotic plankton using metabarcoding revealed substantial diversity of parasitic dinoflagellates in the order of Syndiniales throughout the oceans [7], and highlighted the top-down interactions of the parasitic dinoflagellates in the Syndiniales and their host organisms [8]. Among these, parasitic dinoflagellates in the genus *Amoebophrya* and members of the family Parviluciferaceae are comparatively well researched, as they infect planktonic dinoflagellates including several harmful algal bloom (HAB) species. Beyond these studies, the current knowledge of the biological characteristics and ecological roles of parasitic dinoflagellates are limited compared to other marine protists in sister phyla (ex. planktonic dinoflagellates, apicomplexan, ciliates).

The parasitic dinoflagellate *Hematodinium* spp. is an endoparasitic dinoflagellate infecting a wide range of marine decapod crustaceans [9]. Since its initial description, it had been reported infecting over 40 species of marine crustaceans, many of which support important commercial fisheries [10–12]. Within the host, the parasite resides and proliferates in hemolymph or haemocoel, causing tissue damage and malfunction of infected organs (ex. heart, hepatopancreas, gills) eventually leading to mortality in the late stages of infection [13, 14]. In the last three decades *Hematodinium* spp. infection have been reported in a number of new hosts and locations, and at higher prevalence in species where the prevalence was traditionally lower [15–23]. Whether this represents a true range expansion of the parasite, or increased awareness, interest and reporting is debatable, however there is no doubt that the list of susceptible species is growing rapidly. In recent years, *Hematodinium perezi* has been reported from several aquacultured decapod and co-cultured shrimp species in China [21,23,24,] and in other crustaceans residing within and adjacent to polyculture ponds [25, 26] and represents a direct threat to the sustainable culture of marine crustaceans in these regions.

The immune system of crustaceans has been extensively studied [27–30]. Particularly, a number of researches on the immune responses of crustacean hosts against viral infections, including white spot syndrome virus [31], yellow head virus [32], infectious hypodermal and haematopoietic necrosis virus [33], decapod iridescent virus [34], etc., as well as typical bacterial pathogens (e.g. *Vibrio parahaemolyticus*, *V. alginolyticus*, *Spiroplasma eriocheiris*) [35–40]. However, the current knowledge on the interaction between parasites and their crustacean hosts is still extremely limited. Crustaceans lack an adaptive immune system and depend primarily on the innate immunity to defend themselves against pathogens. Recent studies indicated that *H. perezi* could affect multiple key immune pathways in *Portunus trituberculatus*, including the proPO-activating pathway, Toll pathway, NO/O_2_^−^ - generating and scavenging pathways [41–46]. In contrast, Rowley et al. (2015) found no evidence regarding the involvement of circulating hemocytes or fixed or free phagocytes in the immune defense of *Cancer pagurus* against *Hematodinium* sp. parasites, and speculated that this species of *Hematodinium* was not effectively recognized as foreign by the host immune system [47]. Similar host-parasite interactions had been observed in a phylogenetic relative – *Plasmodium*, which evade the immune system by mimicking the naturally-expressed host cellular surface proteins or glycans [48–50]. Considering that *Hematodinium* is able to survive and proliferate in the hemolymph or haemocoel of infected crustacean hosts, the parasite may impose immunomodulatory or immune-suppressive effects. The underlying mechanisms of how *Hematodinium* parasites interact with and modulate its host immune system remain to be further explored.

Recent studies on the taxonomy and molecular phylogeny of dinoflagellates, apicomplexans and ciliates in the superphylum Alveolata further support the close phylogenetic relationship of parasitic dinoflagellates with other well-studied protists (e.g. Apicomplexans) [3,51–53], and provide valuable comparative systems that will help resolve the host-parasite interactions between *Hematodinium* and its crustacean hosts. In addition, the application of omics technologies provides a deeper and broader understanding of the host-parasite interaction [54]. In the present study, an integrated omics approach using microRNAs (miRNAs) transcriptomics and an isobaric labelling method for relative or absolute quantitation (iTRAQ) proteomics were employed to study how *H. perezi* parasites interacted with and modulate the immune system of a marine crustacean, the Chinese swimming crab *Portunus trituberculatus*. The results provide fundamental knowledge on the host-parasite interaction of a parasitic dinoflagellate and its crustacean host, and contribute to a greater understanding of crustacean immune response.

## Results

### Development of *Hematodinium perezi* infection in experimentally inoculated crabs

By day 4 post inoculation (p.i.), *H. perezi* parasites were not observed microscopically by the hemolymph smear assay in the experimentally inoculated crabs, while 7 out of 10 crabs were faintly positive for the presence of *Hematodinium* ITS1 rDNA using a diagnostic PCR assay as described in Small et al. (2007a) [55]. By day 8 p.i., 3 of the experimentally inoculated crabs were diagnosed as positive by microscopic observation, with 8 out of 10 crabs further confirmed with *Hematodinium* infection by PCR assay, and the positive samples were classified as having light infections. By day 16 p.i., 9 out of 10 crabs had developed moderate *Hematodinium* infections, with an abundance of 5.18±0.96×10^5^ parasite cells ml^-1^ in the hemolymph calculated by a hemacytometer. Assessment of the abundance of *Hematodinium* parasites in the hemolymph of infected crabs revealed a gradual progress of infections over time (day 4, 1.14±0.15×10^7^ copies ml^-1^; day 8, 3.29±0.78×10^7^ copies ml^-1^; day 16, 1.85±0.54×10^8^ copies ml^-1^). No experimental crabs died during the experiment. Six out of thirty crabs sampled at day 4, 8 and 16 p.i. were diagnosed as negative by both the macroscopic and standard PCR assays. The reasons for this are unclear but may be due to low abundance of the parasite in the hemolymph below the detection limit of the PCR assay (0.3 parasites per 100 ml hemolymph, Small et al., 2007a) [55] or the presence of uninfected immune crabs as described in Shields & Squyars (2000) [56].

### Hemocyte MiRNAs transcriptomic responses to *H. perezi* parasites

The length distribution analysis showed that the majority of unique miRNA sequences were distributed predominantly in the 21-23 nucleotides (nts) range, and that 22-nt sequences were the most abundant (S1 Fig). After being further processed as Li et al. (2018) [45], a total of 153 miRNAs were identified from the hemocyte samples collected at different time points (4, 8 and 16 days p.i.) after *Hematodinium* challenge (S2 Fig). A total of 44 differentially expressed miRNAs (DEMs) were identified and showed significant activation or suppression in hemocytes populations (Fig 1). Compared to the transcripts obtained in the control samples (day 0), 20 DEMs (17 up- regulated, 3 down-regulated) were identified in hemocytes collected 4 days post inoculation (dpi), 8 DEMs (6 up-regulated, 2 down-regulated) were identified in hemocytes collected at 8 dpi and 19 DEMs (14 up-regulated, 5 down-regulated) were identified in hemocytes collected at 16 dpi (Fig 2).

**Fig 1.**
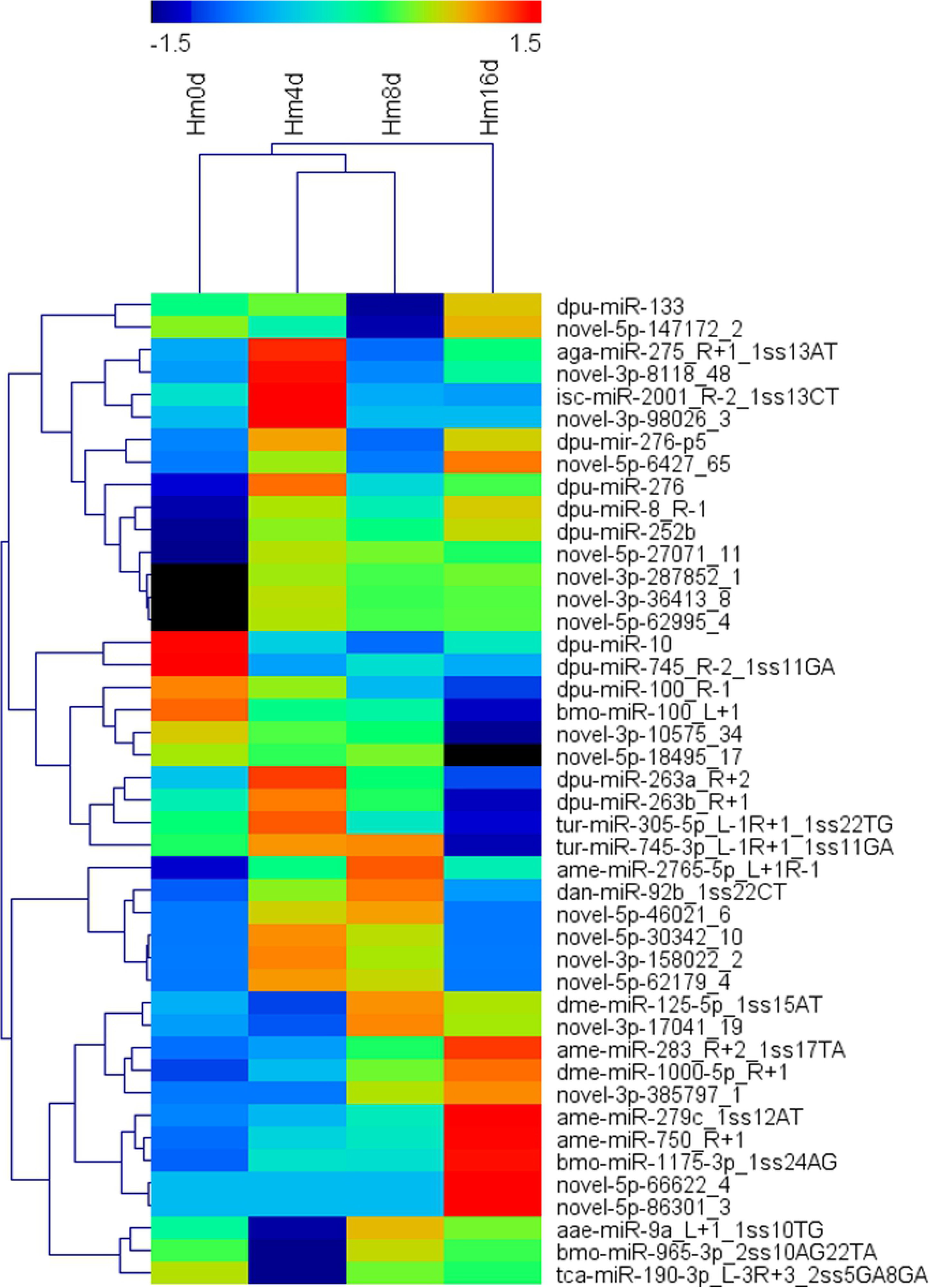
Heat maps of the differentially expressed miRNAs (DEMs). Hierarchical clustering of the 44 significantly changed miRNAs in hemocytes at 4, 8 and 16 days post challenge with *Hematodinium perezi*. Colors represent higher or lower expression levels relative to the controls.

**Fig 2.**
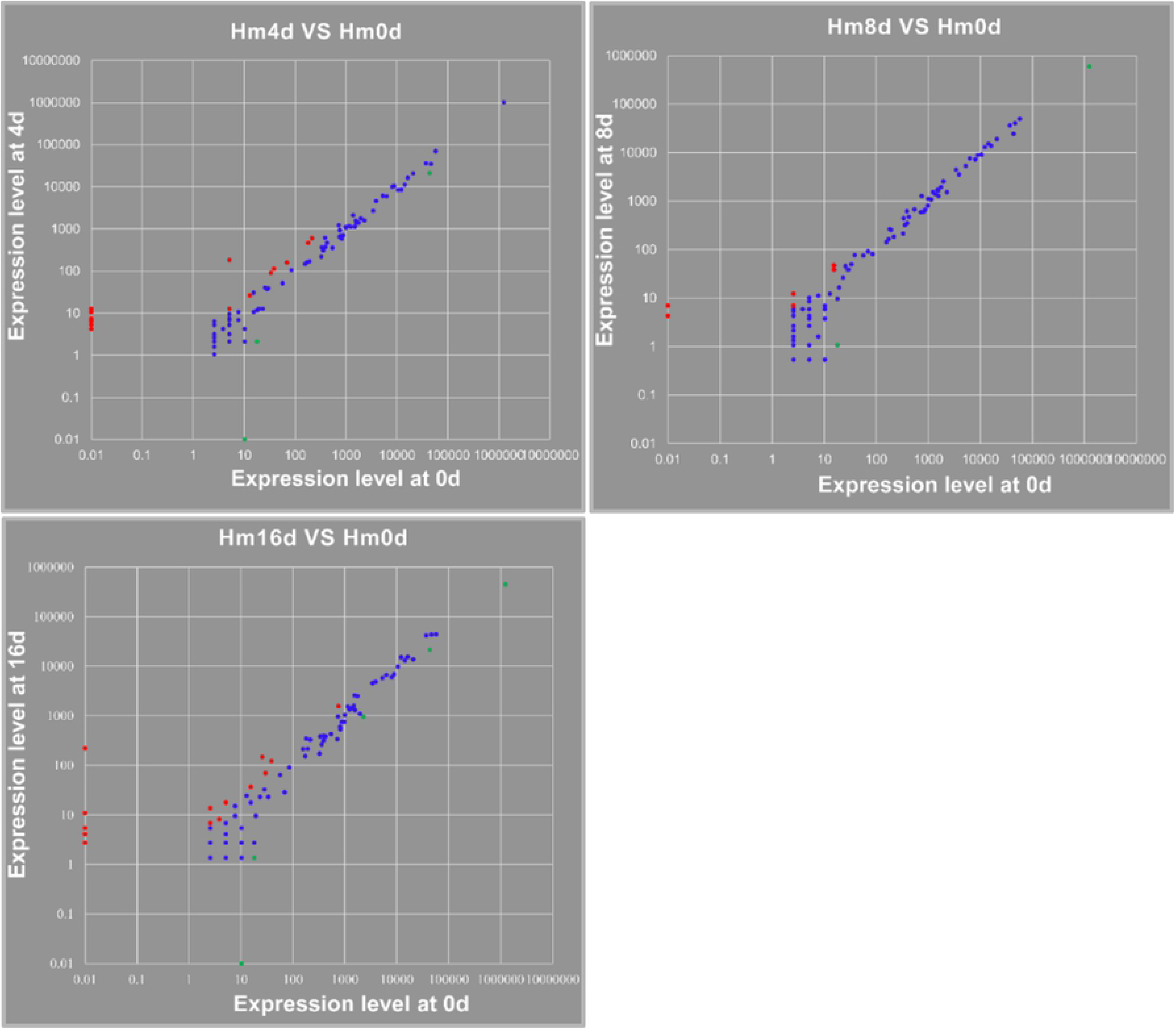
Expression analysis of the identified miRNAs. The expression levels of the miRNAs were detected at different time points of 4 d (Hm4d VS Hm0d), 8 d (Hm8d VS Hm0d) and 16 d (Hm16d VS Hm0d) post challenge with *H. perezi* and compared to the time point 0 d (Hm0d). The red dots represent the significantly up-regulated miRNAs, while the green dots represent the significantly down-regulated ones. The blue dots represent the non-significantly changed ones.

After assessed with the Targetscan 6.2 and miRanda algorithms, a total of 964 target genes of the 44 DEMs were predicted. Kyoto Encyclopedia of Genes and Genomes (KEGG) pathway analysis of the target genes indicated that 34 immune related pathways, including typical cellular (e.g. Apoptosis, Endocytosis, etc.) and humoral immune pathways (e.g. Toll-like receptor signaling pathway, Jak-STAT signalling pathway, complement and coagulation cascades, etc.), were modulated post-transcriptionally by *Hematodinium* parasites (Fig 3).

**Fig 3.**
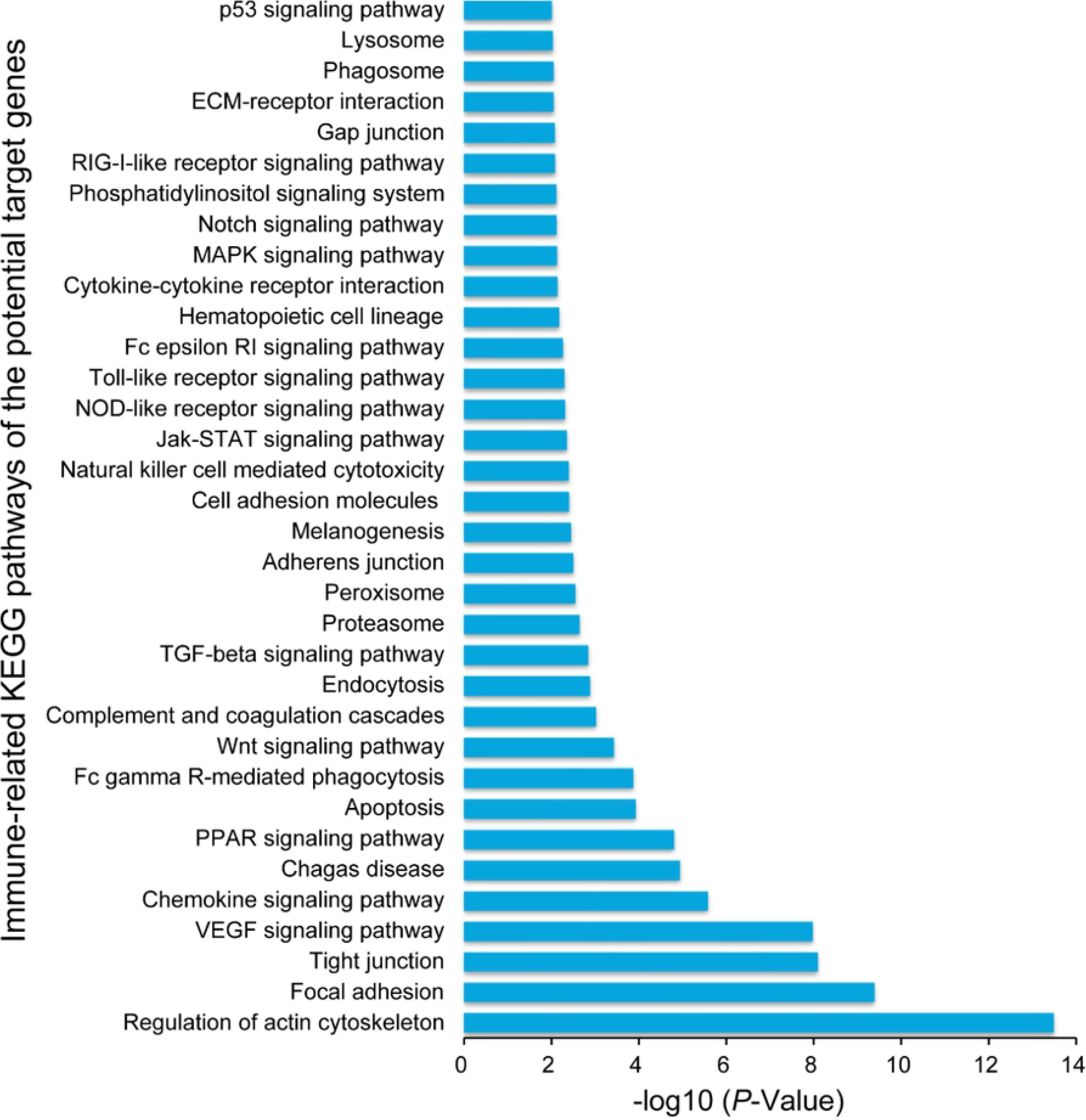
The significantly enriched immune-related KEGG pathways of the predicted target genes of the DEMs.

### Hemocyte proteomic expression to *H. perezi* parasites

Obvious proteomic alterations were detected using the iTRAQ technique in hemocytes of *P. trituberculatus* challenged with *H. perezi*. A total of 1977 proteins were identified in the hemocytes samples and classified into six categories of KEGG pathways (e.g. Organismal Systems, Metabolism, Cellular Processes, etc.), including 523 differentially expressed proteins (DEPs) (S3 Fig, Fig 4). The proteomic expression of these DEPs varied in the hemocytes samples collected at the different time points post *Hematodinium* parasite challenge. Compared to the control samples (day 0), there were 155 proteins up-regulated and 123 proteins down-regulated at 4 dpi, 66 up- regulated proteins and 152 down-regulated proteins at 8 dpi, 71 up-regulated proteins and 128 down-regulated proteins at 16 dpi (Fig 5). Upon KEGG pathway analysis of all the DEPs, 22 immune related pathways (Table 1) were found to be modulated by *Hematodinium* parasites.

**Fig 4.**
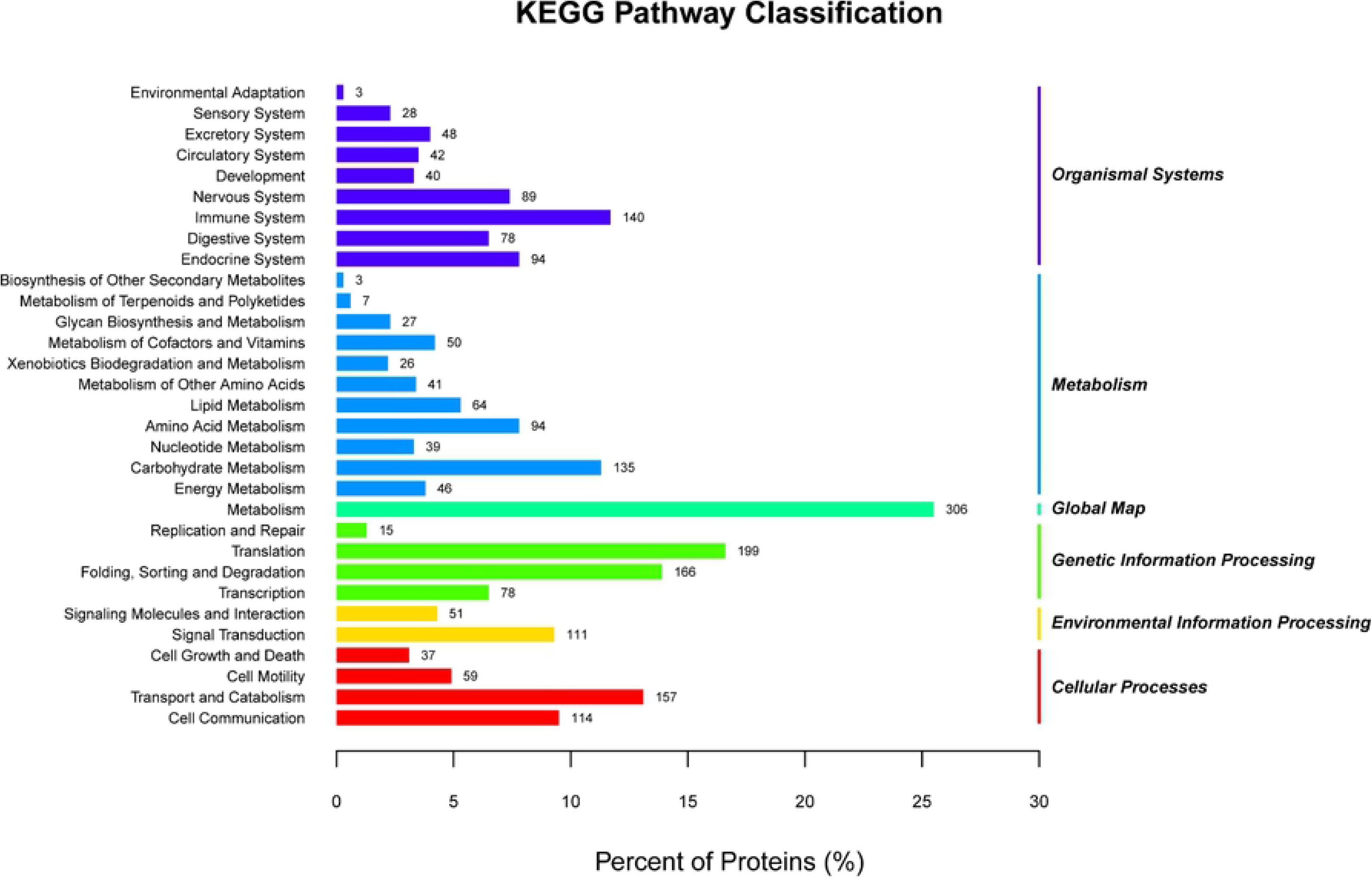
KEGG pathway classification of the identified proteins in hemocytes.

**Fig 5.**
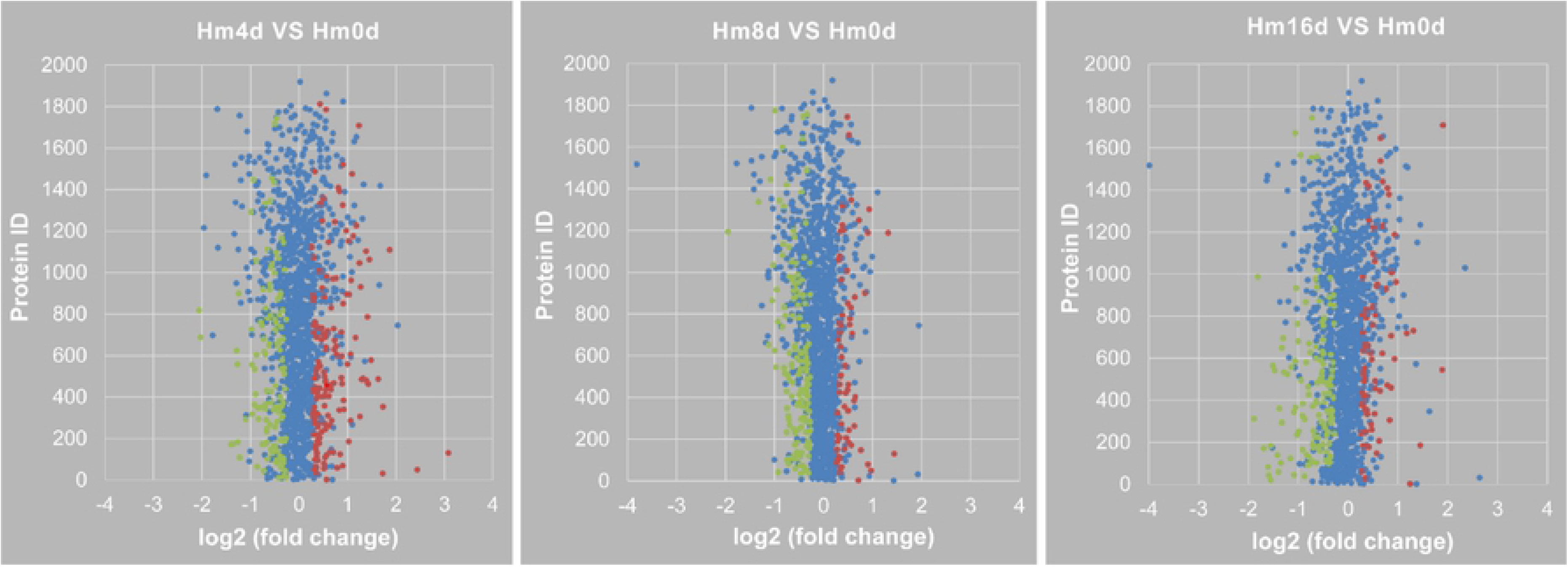
Expression analysis of the identified proteins. Hemocyte protein levels were detected at different time points of 4 d (Hm4d), 8 d (Hm8d) and 16 d (Hm16d) post challenge with challenge with *H. perezi* and compared to the time point 0 d (Hm0d, control). The red dots represent the significantly up-regulated proteins, while the green dots represent the significantly down-regulated proteins. The blue dots represent the non-significantly changed ones.

**Table 1.**
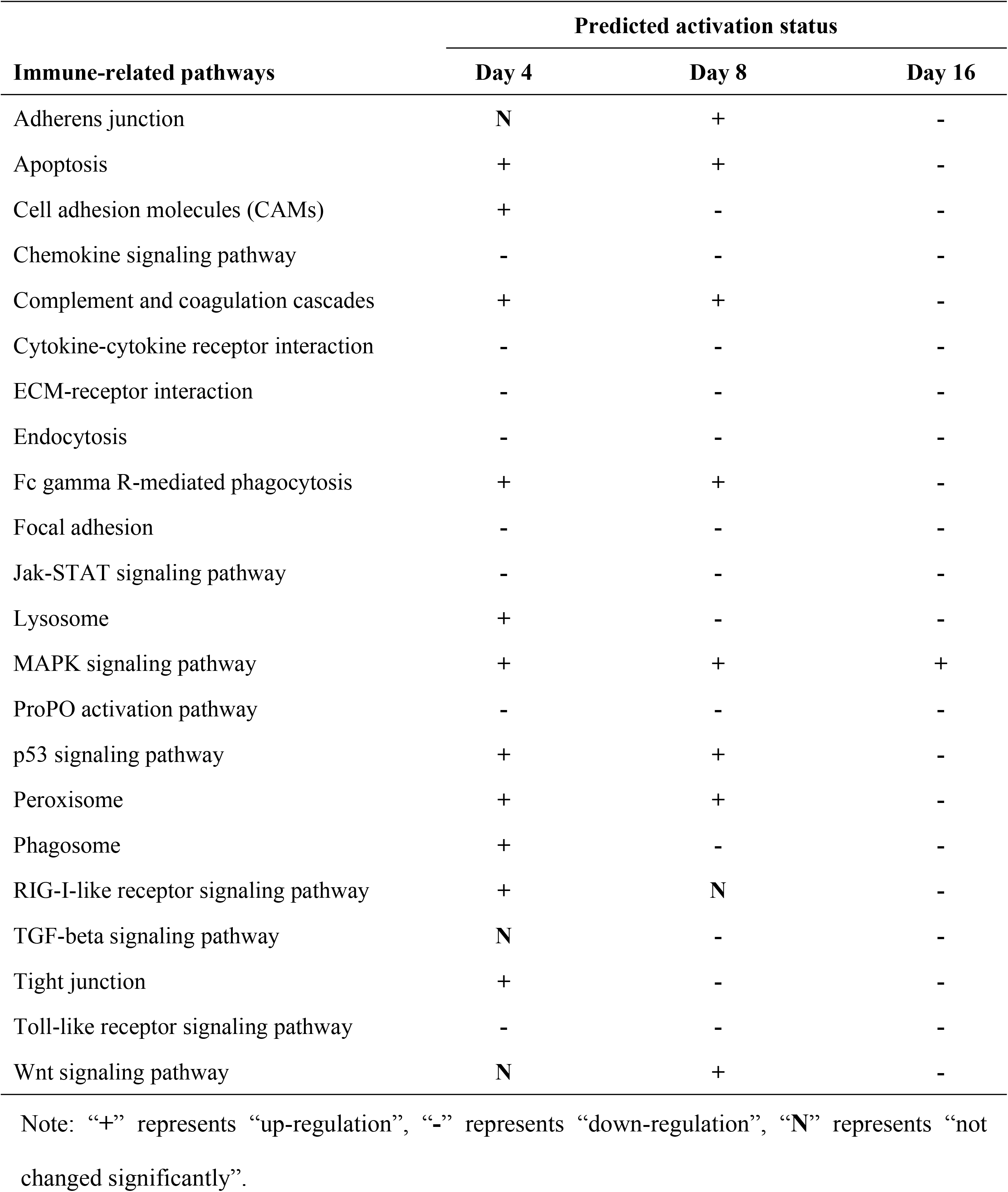
The predicted activation status of the significantly enriched immune-related pathways at different time points after challenge with *Hematodinium perezi*.

In addition, the activation status of each significantly enriched immune-related pathway (Table 1) was estimated based on the alteration and biological function of the DEPs. Due to the difference in the degree of *Hematodinium* infection, the hosts’ immune responses to the parasitic infection was different at various time points. Particularly, the critical immune-related pathways (e.g. Toll-like receptor signaling pathway, Jak-STAT signaling pathway, Chemokine signaling pathway, Endocytosis, Cytokine-cytokine receptor interaction, proPO activation pathway and TGF-beta signaling pathway) were down-regulated in hemocytes from crabs inoculated with *Hematodinium* parasites. The pathways involved in the hosts’ cellular homeostasis (e.g. Phagosome, Peroxisome, Lysosome) were up-regulated in crabs with vary light or light infection (at 4 dpi and 8 dpi), while down-regulated in hemocytes from crabs with moderate infection (at 16 dpi). The pathways associated with hosts’ cellular communication and structure (e.g. ECM-receptor interaction, Focal adhesion, Tight junction, Cell adhesion molecules) were mostly downregulated in hemocytes from crabs with light and moderate infections (at 8 dpi and 16 dpi). The pathways of Apoptosis, p53 signaling pathway and Complement and coagulation cascades were up- regulated in hemocytes from crabs with very light or light infections (at 4 dpi and 8 dpi) and down-regulated in the hemocytes from crabs with moderate infection (at 16 dpi). The pathways of Adherens junction, Wnt signaling pathway, Fc gamma R-mediated phagocytosis and RIG-I-like receptor signaling pathway were up-regulated in the hemocytes from crabs with very light or light infection (at 4 dpi and 8 dpi), but were down-regulated in the hemocytes from crabs with moderate infection (at 16 dpi). The MAPK signaling pathway was up-regulated in the hemocytes from all crabs after challenged by *Hematodinium* parasites.

### Concordance of the immune-related KEGG pathways associated with *H. perezi* infection

By comparing the immune related pathways enriched from the potential target genes of the DEMs with those enriched from the DEPs, 22 immune related pathways were common in hemocytes samples collected from *P. trituberculatus* crabs challenged with *H. perezi*. With regard to the common immune-related pathways mentioned above, the target genes of the DEMs were used to search the proteomics data. Thirteen significantly changed proteins were identified as the candidate miRNA targets of 12 DEMs and mainly involved in modulating 9 critical immune related pathways (Table 2). The pathways of ECM-receptor interaction, focal adhesion, nodulation and antimicrobial peptides were downregulated in hemocytes from all crabs challenged with *Hematodinium* parasites. The pathways of phagosome, peroxisome and lysosome were up-regulated in hemocytes from crabs with very light infection (at 4 dpi) while down-regulated in hemocytes from crabs with moderate infections (at 16 dpi). The complement and coagulation cascades pathway was up-regulated in hemocytes from crabs with light infections (at 8 dpi). The MAPK signaling pathway was up-regulated in all hemocytes from crabs challenged with *Hematodinium* parasites.

**Table 2.**
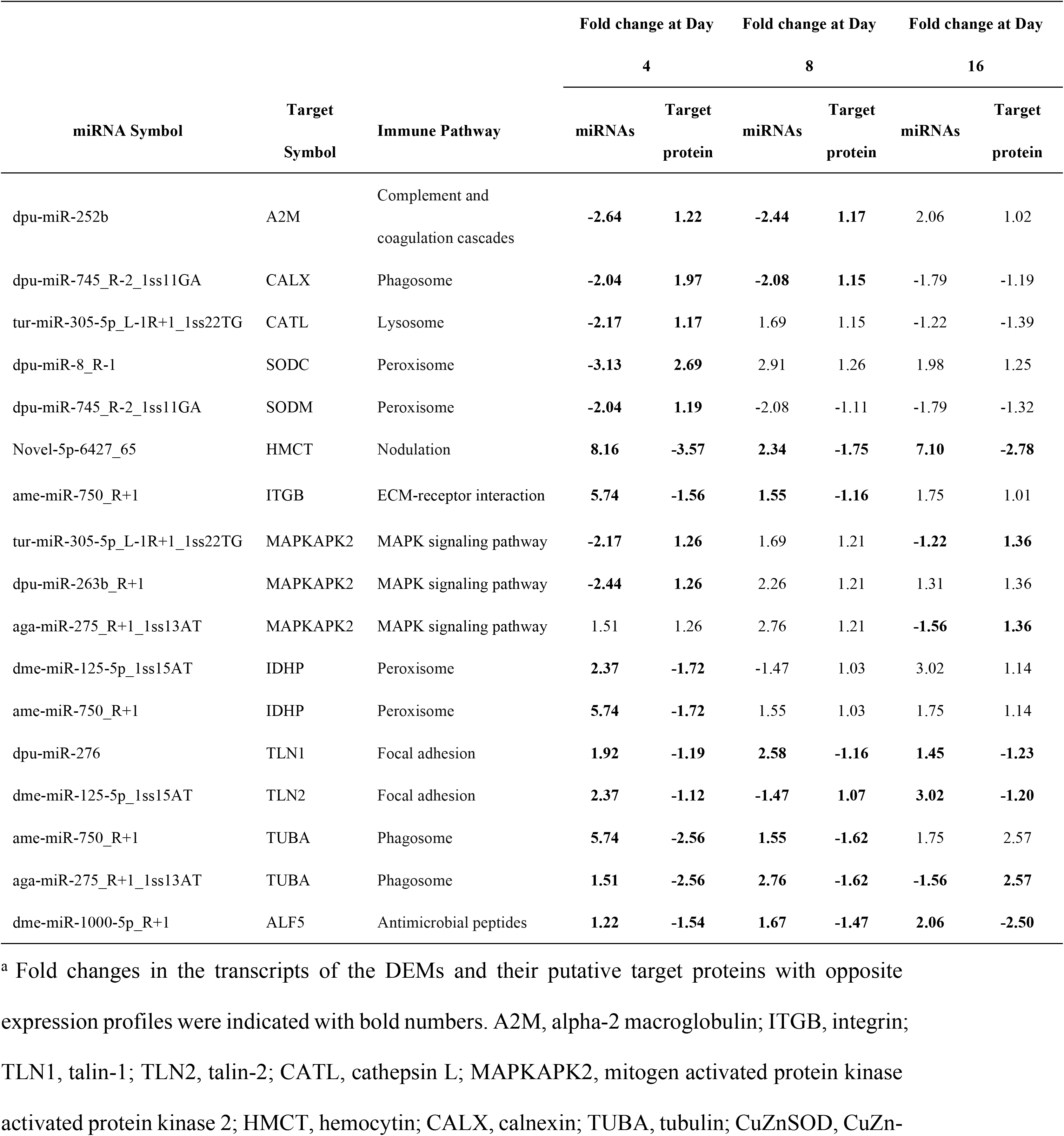
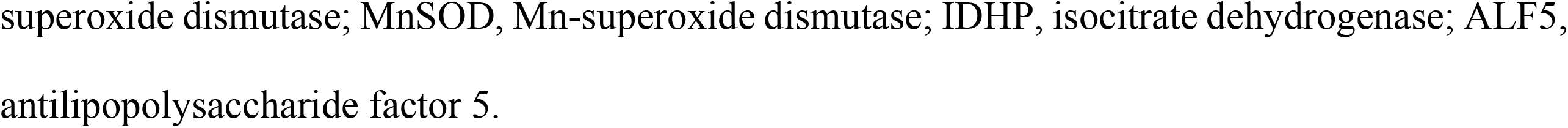
Variation in the expression of the identified differentially expressed miRNAs (DEMs) and their target proteins involved in the immune-related pathways^a^.

| miRNA Symbol | Target Symbol | Immune Pathway | Fold change at Day 4 |  | Fold change at Day 8 |  | Fold change at Day 16 |  |
| --- | --- | --- | --- | --- | --- | --- | --- | --- |
|  |  |  | miRNAs | Target protein | miRNAs | Target protein | miRNAs | Target protein |
| dpu-miR-252b | A2M | Complement and coagulation cascades | **-2.64** | **1.22** | **-2.44** | **1.17** | 2.06 | 1.02 |
| dpu-miR-745_R-2_1ss11GA | CALX | Phagosome | **-2.04** | **1.97** | **-2.08** | **1.15** | -1.79 | -1.19 |
| tur-miR-305-5p_L-1R+1_1ss22TG | CATL | Lysosome | **-2.17** | **1.17** | 1.69 | 1.15 | -1.22 | -1.39 |
| dpu-miR-8_R-1 | SODC | Peroxisome | **-3.13** | **2.69** | 2.91 | 1.26 | 1.98 | 1.25 |
| dpu-miR-745_R-2_1ss11GA | SODM | Peroxisome | **-2.04** | **1.19** | -2.08 | -1.11 | -1.79 | -1.32 |
| Novel-5p-6427_65 | HMCT | Nodulation | **8.16** | **-3.57** | **2.34** | **-1.75** | **7.10** | **-2.78** |
| ame-miR-750_R+1 | ITGB | ECM-receptor interaction | **5.74** | **-1.56** | **1.55** | **-1.16** | 1.75 | 1.01 |
| tur-miR-305-5p_L-1R+1_1ss22TG | MAPKAPK2 | MAPK signaling pathway | **-2.17** | **1.26** | 1.69 | 1.21 | **-1.22** | **1.36** |
| dpu-miR-263b_R+1 | MAPKAPK2 | MAPK signaling pathway | **-2.44** | **1.26** | 2.26 | 1.21 | 1.31 | 1.36 |
| aga-miR-275_R+1_1ss13AT | MAPKAPK2 | MAPK signaling pathway | 1.51 | 1.26 | 2.76 | 1.21 | **-1.56** | **1.36** |
| dme-miR-125-5p_1ss15AT | IDHP | Peroxisome | **2.37** | **-1.72** | -1.47 | 1.03 | 3.02 | 1.14 |
| ame-miR-750_R+1 | IDHP | Peroxisome | **5.74** | **-1.72** | 1.55 | 1.03 | 1.75 | 1.14 |
| dpu-miR-276 | TLN1 | Focal adhesion | **1.92** | **-1.19** | **2.58** | **-1.16** | **1.45** | **-1.23** |
| dme-miR-125-5p_1ss15AT | TLN2 | Focal adhesion | **2.37** | **-1.12** | **-1.47** | **1.07** | **3.02** | **-1.20** |
| ame-miR-750_R+1 | TUBA | Phagosome | **5.74** | **-2.56** | **1.55** | **-1.62** | 1.75 | 2.57 |
| aga-miR-275_R+1_1ss13AT | TUBA | Phagosome | **1.51** | **-2.56** | **2.76** | **-1.62** | **-1.56** | **2.57** |
| dme-miR-1000-5p_R+1 | ALF5 | Antimicrobial peptides | **1.22** | **-1.54** | **1.67** | **-1.47** | **2.06** | **-2.50** |

<sup>a</sup> Fold changes in the transcripts of the DEMs and their putative target proteins with opposite expression profiles were indicated with bold numbers. A2M, alpha-2 macroglobulin; ITGB, integrin; TLN1, talin-1; TLN2, talin-2; CATL, cathepsin L; MAPKAPK2, mitogen activated protein kinase 2; HMCT, hemocytin; CALX, calnexin; TUBA, tubulin; CuZnSOD, CuZn-
superoxide dismutase; MnSOD, Mn-superoxide dismutase; IDHP, isocitrate dehydrogenase; ALF5, antilipopolysaccharide factor 5.

### Validation of omics analysis by qRT-PCR

The transcripts of ten immune-related DEMs were detected by qRT-PCR assays to validate the miRNA transcriptomic results (Fig 6). The transcripts of aga-miR- 275_R+1_1ss13AT were increased significantly at 4 dpi and 8 dpi, then decreased at 16 dpi. The transcripts of ame-miR-750_R+1 were increased significantly at 4 dpi. The transcripts of dme-miR-1000-5p_R+1, dpu-miR-276 and dpu-miR-252b were increased significantly during the whole period of *Hematodinium* challenge. The transcripts of dme-miR-125-5p_1ss15AT were increased significantly at 4 dpi and 16 dpi. The transcripts of dpu-miR-263b_R+1 and dpu-miR-8_R-1were initially decreased significantly at 4 dpi, then increased significantly at 8 dpi and 16 dpi. The transcripts of dpu-miR-745_R-2_1ss11GA were decreased significantly during the whole period of *Hematodinium* challenge. The transcripts of tur-miR-305-5p_L-1R+1_1ss22TG were initially decreased significantly at 4 dpi, then increased at 8 dpi. All qRT-PCR results showed a consistency with the transcriptome data and provided further support for the reliability of the small RNA sequencing data.

**Fig 6.**
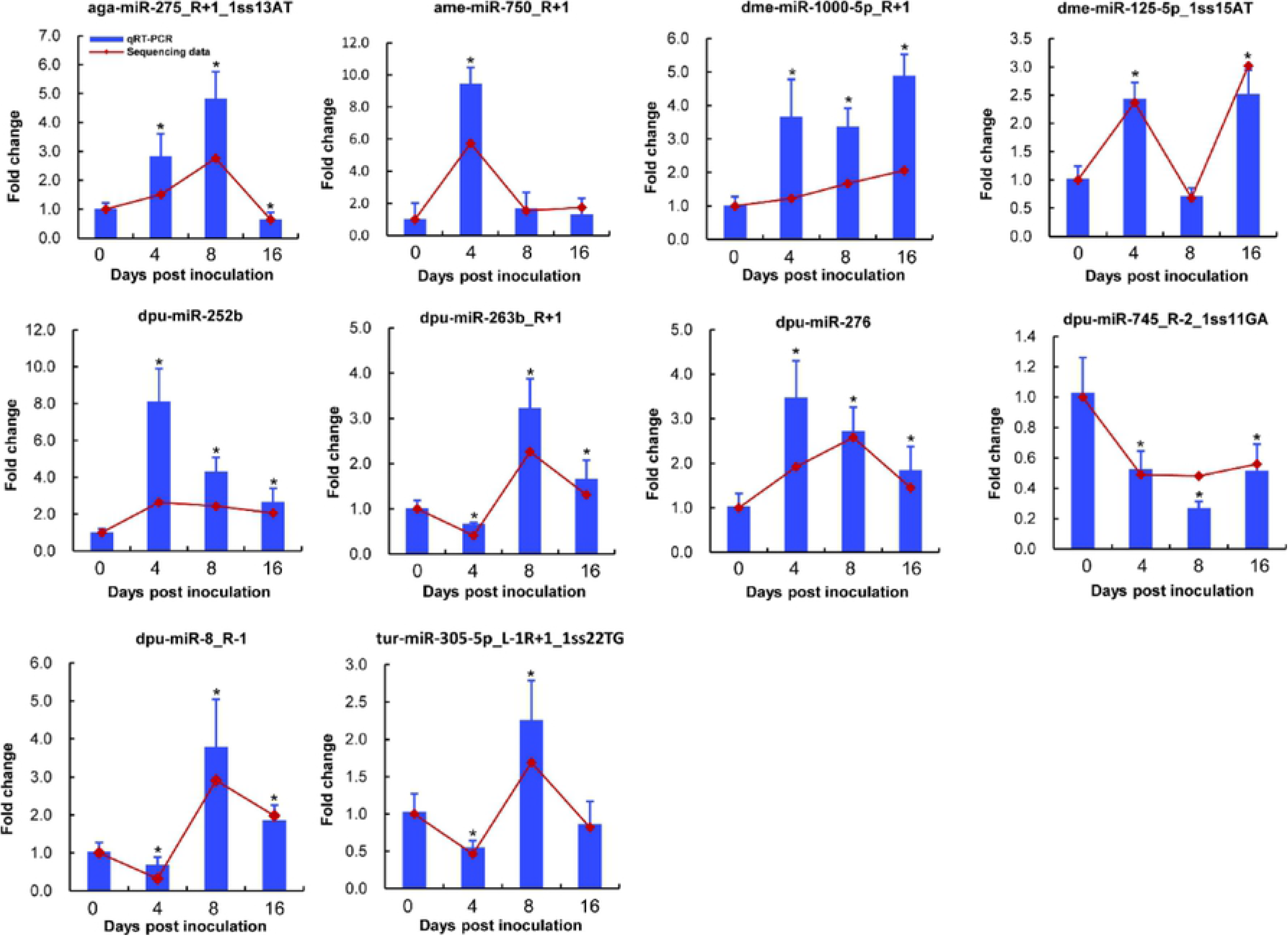
Validation of 10 DEMs by qRT-PCR. The qRT-PCR results and the sequencing data are represented with the bar and line chart, respectively. The qRT-PCR results with statistical significances are indicated with asterisks compared to the time point of 0 d (*P* < 0.05).

To further confirm the iTRAQ results, the mRNA transcripts of ten DEPs which were involved in key immune-related pathways including MAPK signaling pathway, Lysosome, Complement and coagulation cascades, Phagosome, ECM-receptor interaction, and Peroxisome were also evaluated by qRT-PCR assays (Fig 7). The transcripts of *CuZnSOD*, *MnSOD*, *TALIN1* and *MAPKAPK2* were significantly increased at 4 dpi or/and 16dpi. The transcripts of *cathepsin L* were significantly increased at 4 dpi and 8 dpi, in which time the transcripts of *Α2M* were significantly decreased. While, the transcripts of *hemocytin* were significantly decreased in all crabs post *Hematodinium* challenge. The transcripts of *α-spectrin* and *β-integrin* were depressed significantly at 4 dpi, while *β-integrin* enhanced significantly at 16 dpi. The transcripts of *calnexin* were significantly up-regulated at 4 dpi, while down-regulated significantly at 16 dpi. Overall, the qRT-PCR results were consistent with the iTRAQ-based proteomics data, reflecting the reliability of the iTRAQ-based proteomics results in the present study.

**Fig 7.**
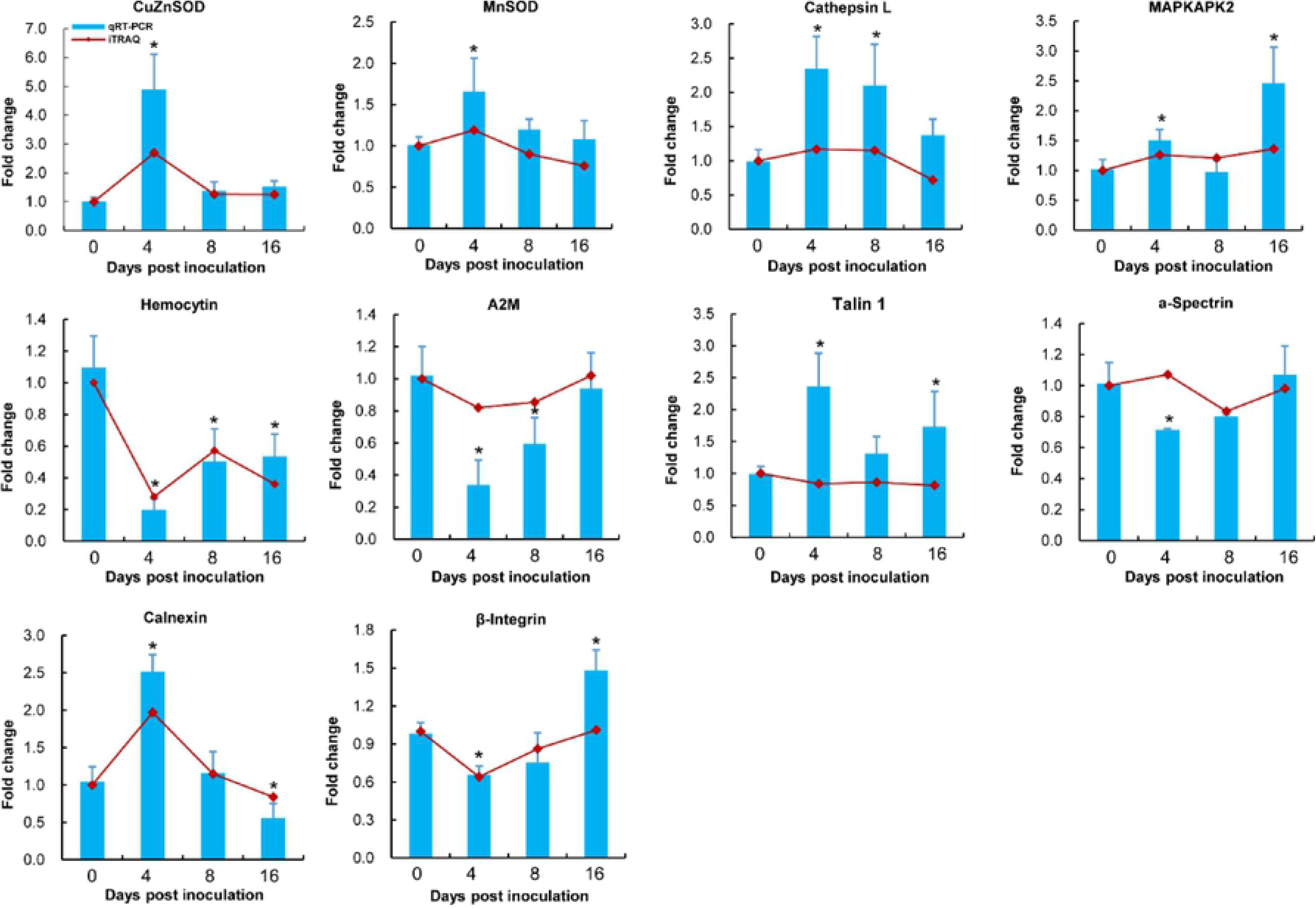
Comparison of transcriptional analysis and iTRAQ-based proteomic results of ten DEPs. The fold changes in transcripts of the ten DEPs in hemocytes of *P. trituberculatus* challenged with *H. perezi* were assayed by qRT-PCR. The qRT-PCR and the iTRAQ-based proteomic results are represented with the bar and line chart, respectively. Transcriptional changes with statistical significances are indicated with asterisks compared to the time point of 0 d (*P* < 0.05).

## Discussion

The parasitic dinoflagellate *Hematodinium* is able to survive and proliferate in the hemolymph or haemocoel of its crustacean host, eventually leading to mortality of the infected host due to organ dysfunction [13]. The suppressive effect of the parasite on the host crustaceans immunity has previously been observed in the early stage of *Hematodinium perezi* infections in *Portunus trituberculatus* [41–46], and potential parasite immune evasion has been speculated during the parasite-host interaction process [47]. The present integrative omics approach in the Chinese swimming crab *P. trituberculatus*, further revealed the immunomodulatory effects of *H. perezi* on the host crustacean immunity over time, in which the parasitic dinoflagellate caused significant miRNome modulation and proteome reorganization in hemocytes of *P. trituberculatus*. Particularly, *H. perezi* affected cellular immunity by damaging the intercellular communication and integrity of hemocytes, caused immunomodulatory effects on humoral immunity by inhibiting critical pattern recognition receptors (PRRs)-mediated immune responses, and triggering cellular oxidative stress via imbalance of host’s redox homeostasis.

Hemocytes are the most important component of the crustacean immune system and play an important role in the cellular and humoral immune response against invading pathogens [27, 57]. *H. perezi* infection of the blue crab *Callinectes sapidus* resulted in a decrease of hemocyte density, loss of clotting ability and the alteration of hemolymph proteins or glycogen levels, and caused eventual mortality of infected hosts [56, 58]. In addition, a host nodulation response of hemocytes to *Hematodinium* parasites was observed in several infected crustacean species [59, 60]. In the present study, *H. perezi* parasites overtly modulated the intercellular communication and architecture of hemocytes in *P. trituberculatus* by affecting the following pathways “Cell adhesion molecules”, “ECM-receptor interaction”, “Focal adhesion”, “Adherens junction” and “Tight junction”. Cell adhesion molecules (CAMs) play important roles in maintaining tissue integrity and mediating migration of immune cells [61, 62], which are involved in invertebrate cellular immunity such as phagocytosis and nodulation [63, 64]. Nodulation is a prompt response to eliminate invading microorganisms in the haemocoel, in which hemocytin plays important role in hemocyte aggregation and nodule formation [65]. The significant inhibition of hemocytin in hemocytes of *P. trituberculatus* challenged with *H. perezi* may therefore contribute to the loss of clotting capability reported to occur in the hemolymph of infected hosts [22,25,58], thereby facilitating parasite escape from the host nodulation response.

The extra cellular matrix (ECM) is an important structure essential for normal tissue homeostasis [66]. ECM components also act as ligands to transmit signals to regulate cellular activities (e.g. cellular adhesion, apoptosis, proliferation, survival) [67]. Dysregulation of ECM components has been previously reported being associated with pathological conditions and disease progression [66], and could be remodeled by the bacterial infection [68]. Recent studies also showed that ECM components (e.g. clotting protein, matrix metalloproteinase-14) were involved in the innate immunity and hematopoiesis of crustacean hosts [69, 70]. In the hemocytes of *P. trituberculatus* challenged with *H. perezi*, several key ECM components, including integrin, laminin, collagen and tenascin, were overtly dysregulated at protein levels, which was observed in the iTRAQ proteomics analysis (unpublished data). In addition, cellular junctions are crucial for tissue structural integrity and pathogen defense [71, 72], and the crucial intercellular junctions (e.g. adherens junction, tight junction) could be disrupted by bacterial or viral infections, benefiting entry or survival of the invading pathogens in vertebrate hosts [73–75], as well as in the crustacean host [64]. In *P. trituberculatus* challenged with *H. perezi*, the pathways of adherens junction and tight junction were also disrupted, with up-regulation in crabs with light infections and down-regulation in crabs with moderate infections. Meanwhile, the cellular immune-related pathways of Endocytosis and ECM-receptor interaction were down-regulated in hemocytes of *P. trituberculatus* challenged with *H. perezi*. Consequently, disruption of intercellular communication and structure in the infected crustacean hosts likely impaired the host cellular immune response, which then contributed to the survival and pathogenesis of *Hematodinium* in its crustacean host.

In the present study, the major humoral immune pathways associated with hemocytes of crustacean hosts were also affected by *H. perezi*. Important humoral immune pathways, such as “Toll-like receptor signaling pathway”, “Jak-STAT signaling pathway”, “proPO activation pathway”, and “RIG-I-like receptor (RLR) signaling pathways”, were significantly down-regulated in hemocytes of *P. trituberculatus* challenged with *H. perezi* parasites. Conversely, these four key immune pathways are generally stimulated by invading bacterial and viral pathogens [29, 76]. Our previous studies had demonstrated that the transcripts of Toll, β-1, 3-glucan binding protein (LGBP) and proPO-activating factor (PPAF) could be significantly inhibited in hemocytes and hepatopancreas of *P. trituberculatus* challenged with *H. perezi*, implying an parasite-driven immunosuppressive mechanism against the host Toll signaling pathway and proPO systems [41,42,44]. The iTRAQ-based proteomics study further suggested that *H. perezi* parasites caused immune-suppressive effects on immune responses of hepatopancreas in *P. trituberculatus* by inhibiting important pattern recognition receptors (e.g. Toll, C-lectin, and SR class B) [46]. Interestingly, important antimicrobial peptides such as anti-lipopolysaccharide factors (ALF3, ALF5, ALF6 and ALF7) were significantly down-regulated in the hemocytes from *P. trituberculatus* challenged with *H. perezi* parasites in the present study. In previous studies, the important humoral immune molecules ALFs were generally up-regulated in response to the bacterial, fungal and viral challenge [77–79], indicating that the inhibitory effects induced by *H. perezi* are likely an important immunomodulatory strategy and worthy of further study.

The health status and immune defense of crustaceans is closely correlated with their cellular redox status [37], and maintenance of the cellular redox homeostasis, especially the levels of the reactive oxygen/nitrogen free radicals (ROS/RNS), is crucial for the normal physiological functions (e.g. cell proliferation, apoptosis, signal transduction) of major organs [80–82]. *H. perezi* was previously shown to induce an imbalance in the redox homeostasis of *P. trituberculatus* via dysregulation of the host cellular ROS/RNS-generating/scavenging system [44]. In the present study, two critical cytosolic superoxide dismutase (MnSOD and CuZnSOD) were significantly upregulated in hemocytes of *P. trituberculatus* challenged with *H. perezi*, further implying the increased level of ROS and the imbalance of host redox homeostasis. Furthermore, the peroxisomes pathway, closely associated with maintenance of the cellular redox homeostasis [83] and the antiviral innate immunity [84, 85], was significantly up-regulated in light *H. perezi* infections. The serious imbalance of cellular redox homeostasis may result in cellular apoptosis and the dysfunction of hemocytes in infected crabs. In addition, the cellular apoptosis-associated signaling pathways “p53 signaling pathway” and “Wnt signaling pathway” in hemocytes of *P. trituberculatus* were significantly up-regulated in light infections. We hypothesize that in light infections the cellular redox imbalance and apoptosis of hemocytes impaired host immunity, and thereby facilitated the successful establishment of *Hematodinium* infections.

Protozoan parasites have evolved a sophisticated epigenetic strategy to regulate the host immune system [86, 87], and ensure their survival by inhibiting host innate defenses through regulation of the host gene expression [88]. Particularly, protozoan parasites can hijack host immune defense functions by subverting the host miRNAs [89]. In *P. trituberculatus* crabs challenged with *H. perezi*, 51 miRNAs were found to be differentially expressed in the hepatopancreas, which were involved in regulating host’s important immune pathways (e.g. Endocytosis, Complement and coagulation cascades, Toll signaling pathway, JAK-STAT signaling pathway, and p53 signaling pathway) [45]. The results from the current study further indicate that *H. perezi* parasites induce post-transcriptional regulatory effects on the immunity of the host crustacean by modulating the host miRNome (Fig 8). The immune-related pathways of phagosome, peroxisome, and lysosome were up-regulated post-transcriptionally at the early stage of infection, which then caused an imbalance of cellular homeostasis. The immune responses of nodulation and production of antimicrobial peptides could be significantly down-regulated via post-transcriptional suppression of the protein expressions of hemocytin and the critical antimicrobial molecule ALF5. In crabs challenged with *H. perezi*, the important proPO activation pathway was significantly suppressed via post-transcriptional up-regulation of the protein expression of A2M, which has been proved to be an important inhibitor on crustacean proPO system [90, 91]. In addition, ECM-receptor interaction and focal adhesion were also down-regulated post-transcriptionally, which would affect cellular structure and communication. Furthermore, the MAPK signaling pathway was up-regulated via post-transcriptional up-regulation of the MAPKAPK2 protein levels. MAPKAPK2 played a vital role in regulating the production of inflammatory cytokine tumor necrosis factor (TNF) [92], which participated in regulating the expression of important antimicrobial molecules (e.g. ALFs, crustins) in crustacean antibacterial immune responses [93–95]. The elevated MAPKAPK2 levels induced by *H*. *perezi* might regulate the production of specific host’s immune effector molecules (e.g. TNFs, ALFs), thereby facilitating the immune-evasion process and survival of the parasites in crustacean hosts. These epigenetic strategies likely help to regulate and evade the host immune system facilitating the survival and proliferation of *Hematodinium* parasites within its crustacean host.

**Fig 8.**
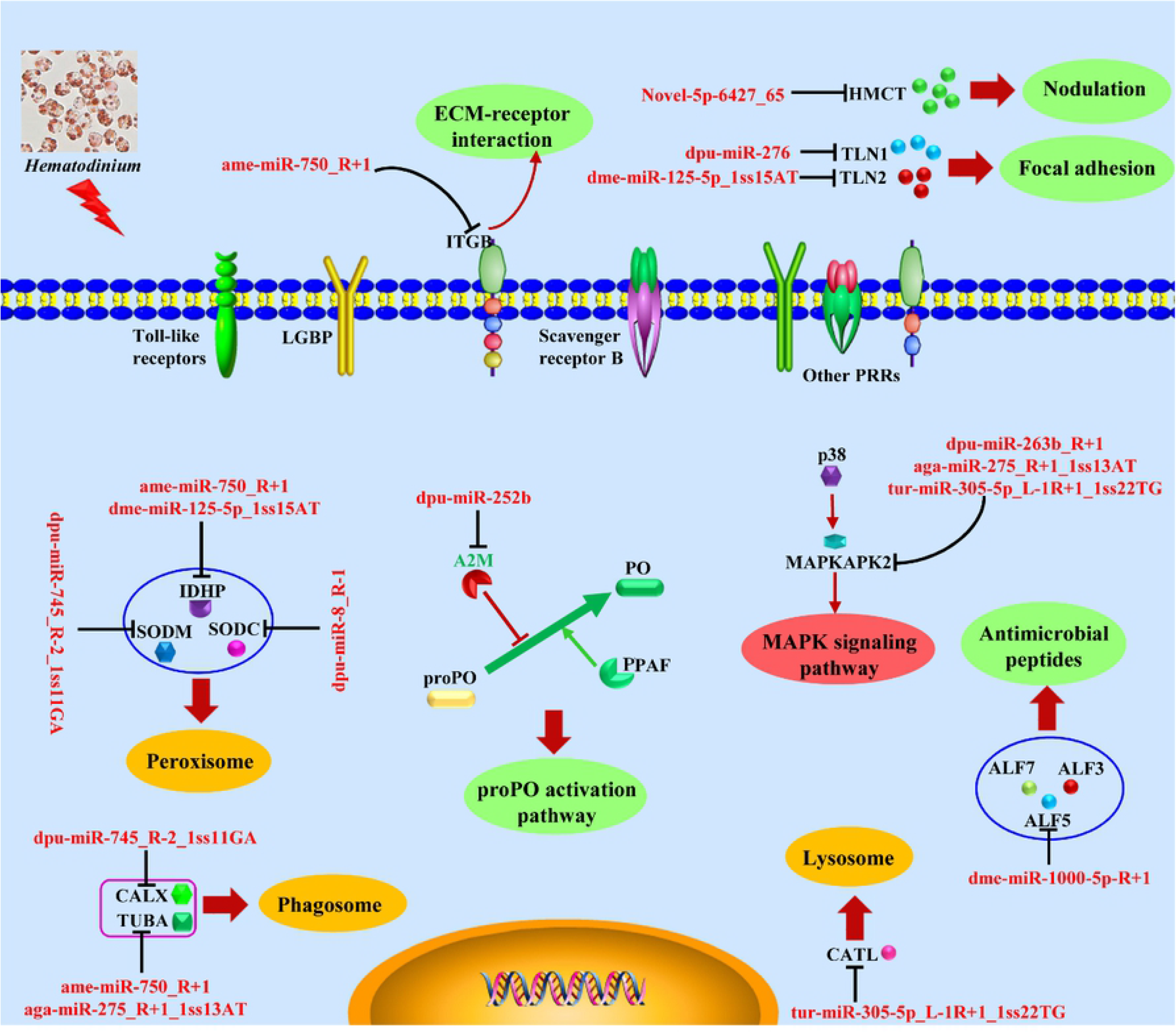
Post-transcriptional regulatory effects of *H. perezi* on immune responses in hemocytes from *P. trituberculatus* by regulating host miRNAs. The pathways of ECM-receptor interaction, focal adhesion, nodulation and antimicrobial peptides were downregulated post *Hematodinium* inoculation. The pathways of phagosome, peroxisome, lysosome and the complement and coagulation cascades were up-regulated at the stage of light infection while down-regulated at the stage of moderate infection. The MAPK signaling pathway was up-regulated after challenge with *H. perezi* whereas the proPO activation pathway was significantly suppressed after challenge with *H. perezi*. Note: the up-regulated, down-regulated and fluctuated pathways were shaded in red, green and yellow, respectively.

Finally, immunomodulatory strategies of apicomplexan parasites are often complex and can also be life-stage specific [48,96–98]. Particular life cycle stages (e.g. *Plasmodium* spp. merozoites) are known to modulate the host response by expressing host cellular surface proteins or glycans [49, 50]. Similarly, the parasitic dinoflagellate *Hematodinium*, which is a phylogenetic relative of apicomplexans, has a complex life history [99, 100]. Indeed, there was notable differences in the enzyme activities of acid phosphatase and leucine arylamidase between *Hematodinium* species (e.g. *H. perezi* ex. *Callinectes sapidus* and *Hematodinium* sp. ex. *Nephrops norvegicus*) [101], implying the differences in their virulence, which likely trigger different immunomodulatory responses in crustacean hosts. Besides, Hamilton et al. (2010) observed distinct physiological responses of three crab species to *Hematodinium* sp. infection, indicated there were host-specific responses against this generalist parasite [102]. Therefore, certain species or life cycle stages of *Hematodinium* parasitizing in the species-specific crustacean hosts may also utilize a similar strategy of molecular mimicry to evade the host immune response [47], which needs to be further studied. The host specificity, as well as the variation in virulence and pathogenicity of *Hematodinium* parasites in different host species, should also be considered in future studies of the parasite - host interaction.

## Materials and Methods

### Ethic statement

This study was carried out according to the Guide for Care and Use of Laboratory Animals by Chinese Association for Laboratory Animal Sciences (No. 2011-2). All of the experimental procedures were performed as humane as possible, and all animals were treated so as to minimize suffering during the experiment.

### Experimental infection and sample collection

Chinese swimming crabs *Portunus trituberculatus* (Female; wet weight: 163.2 ± 8.5 g, carapace length: 143.8 ± 5.9 mm, carapace width: 65.7 ± 2.1 mm) were collected from an aquaculture farm pond located in Huangdao, Shandong province, China. The crabs were initially screened for *Hematodinium perezi* infection using the hemolymph smear assay as described in Stentiford & Shields (2005) [13], and further confirmed with a specific PCR assay [55] prior to the challenge experiment. Uninfected crabs were acclimatized in an aerated recycling seawater system (30 ppt, 23 ± 0.5 °C) for two weeks. Additional individuals with *H. perezi* infections were collected simultaneously from a pond in another aquaculture farm with frequent *Hematodinium* epidemics. The *Hematodinium* species infecting *P. trituberculatus* in this study was identified and characterized as in Li et al. (2013) [22], and the severity of *Hematodinium* infection was assessed microscopically as described in Wang et al. (2017) [23]. The crabs with heavy *H. perezi* infections were held temporarily in a separate seawater tank under the same conditions until being used as donors for subsequent challenge experiments described below.

*H. perezi* trophonts were isolated and purified from heavily infected Chinses swimming crabs as described in Small et al. (2007a) [55], resuspended in Nephrops saline [99] and adjusted to final cell density of 1 × 10^6^ parasite cells/ml for use as an experimental inoculum. The juncture between the basis and ischium of the 5th walking leg of each recipient crab was sterilized with 70% ethanol, and 100 μL of the above trophont preparation was injected into this location using a 1-mL syringe and 27-gauge needle as detailed in Shields & Squyars (2000) [56]. Fifty crabs inoculated with *H. perezi* parasites were maintained in the aerated recycling seawater system until being sampled. During the experiment, crabs were fed clam meat at night (every other day) and remaining food residues were then removed the following morning. Tank waters were changed periodically to ensure the water quality was within acceptable limits (ammonia: 0-0.3 ppm, nitrite: 0-0.6 ppm, pH: 7.4-8.2).

At designated time points post inoculation (day 0, 4, 8 and 16), 10 individuals were randomly selected and sacrificed for hemocyte collection. The hemocytes were separated as described in Wang et al. (2012) [103]. Briefly, 1 mL of hemolymph was drawn from each crab using 1 mL sterile syringe and added into 2 mL PCR tubes containing 1 mL of pre-chilled anticoagulant [104], then the samples were centrifuged at 800 g at 4 °C for 10 min to sediment the hemocytes. The hemocytes samples were immediately frozen in liquid nitrogen and stored at −80 °C. Then at each time point, five individual hemocytes samples, selected from the above collected hemocytes samples with affirmative *Hematodinium* infection, were further processed for omics sequencing and analysis.

### Qualitative and quantitative assessment of *H. perezi* in the hemolymph of experimentally inoculated crabs

A hemolymph smear assay [13] was used to qualitatively assess the presence of *H. perezi* within the inoculated crabs. In addition, molecular methods were employed to quantify parasite densities in crab hemolymph and further validate the infection status. An additional 200μl of hemolymph was also sampled and preserved in 800 μl of 95% ethanol for the parasite quantification. Genomic DNA was extracted from the collected hemolymph samples using a TIANamp Marine Animals DNA Kit (Tiangen, China) according to manufacturer’s instructions. A standard PCR assay [55] was then used to confirm the infection status of sampled experimental crabs. The presence of *Hematodinium* parasites in the hemolymph samples of experimentally inoculated crabs was quantitatively assessed using a quantitative real time PCR assay as described in Li et al. (2010) [105]. Infection status by day 4 p.i. was defined as very light (negative microscopic identification but positive PCR detection). Infection status by day 8 p.i. and 16 p.i. was defined as the light and moderate according to the microscopic quantification of parasites in the hemolymph of sampled crabs [23].

### RNA library construction, sequencing and MiRNAs transcriptomics analysis

#### RNA library construction and sequencing

Total RNA was extracted from hemocytes sampled at different timepoints using a *mir*Vana™ miRNA Isolation Kit (Life Technologies, USA) according to the manufacturer’s procedures. Extracted RNA samples were then quantified and qualified using a Bioanalyzer 2100 and RNA 6000 Nano LabChip Kit (Agilent, USA) with RIN number >7.0. One µg of total RNA was used to construct small RNA libraries using the TruSeq Small RNA Sample Prep Kit (Illumina, USA). The resulting library was sequenced with the single-end sequencing (36 bp) using the Illumina HiSeq 2500 sequencer (LC Sciences, USA) following the manufacturer’s instructions.

#### MiRNAs transcriptomics data analysis

After sequencing, raw image data were transformed by base calling into sequence data called raw sequence data or raw reads. Then, a proprietary pipeline script, ACGT101-miR v4.2 (LC Sciences, USA) was used to analyze the sequencing data. Briefly, after the raw reads were extracted from image data, a series of digital filters (LC Sciences, USA) were applied to remove various unmapped sequencing reads. Then, various ‘‘mappings’’ were performed on unique sequences against pre-miRNA (mir) and mature miRNA sequences (miR) released in the miRBase 22.1, or genome based on the public releases of appropriate species.

### Protein preparation, iTRAQ labeling and proteomics analysis

#### Protein preparation and iTRAQ labelling

Proteins were prepared according to the methods as described in Li et al. (2019) [46]. Briefly, frozen hemocyte samples were resuspended in lysis buffer supplemented with protease inhibitor solution and sonicated on ice before centrifugation and precipitation of proteins. Total protein (100 μg) extracted from each sample was digested with Trypsin Gold (Promega, USA) with the ratio of protein:trypsin = 30:1 and incubated at 37 °C for 16 h. The digested peptides were labelled with iTRAQ reagents according to the manufacturer’s protocols (Applied Biosystems, USA). The peptides from the Hemocyte samples collected at four different time points were labelled with four 8-plex iTRAQ reagents (Applied Biosystems, USA).

#### SCX fractionation

The iTRAQ-labeled peptide mixtures were separated by strong cation exchange chromatography (SCX) using a LC-20AB HPLC Pump System (Shimadzu, Japan) as described in Li et al. (2019) [46]. After reconstituting the dried fractions with buffer A (5% acetonitrile [ACN] and 0.1% formic acid [FA]) to a final concentration of 0.5 μg·μL^−1^, 10 μL of samples were loaded on a LC-20AD nanoHPLC (Shimadzu, Japan) by the autosampler onto a 2-cm C18 trap column (inner diameter 200 μm). Then, the peptides were eluted onto a resolving 10-cm analytical C18 column (inner diameter 75 μm) packed in-house. The liquid chromatography gradient consisted of 5% buffer B (95% ACN and 0.1% FA) for 4 min, 2-35% buffer B for 35 min, 60% buffer B for 5 min, 80% buffer B for 2 min, maintenance at 80% buffer B for 4 min, and finally return to 5% buffer B for 1 min.

#### LC–MS/MS analysis

The MS data of peptide-mixtures were acquired by liquid chromatography - tandem mass spectrometry (LC–MS/MS) analysis via a TripleTOF 5600 System (AB Sciex, Canada) fitted with a Nanospray III source (AB Sciex, Canada) and a pulled quartz tip as the emitter (New Objectives, MA). Data were acquired using an ion spray voltage of 2.5 kV, curtain gas of 30 psi, nebulizer gas of 15 psi, and an interface heater temperature of 150 °C. The MS was operated with a reversed-phase of greater than or equal to 30, 000 full widths at half maximum for time-of-flight MS scans. For information-dependent acquisition, survey scans were acquired in 250 ms, and as many as 30 product ion scans were collected if they exceeded a threshold of 120 counts per second (counts/s) and had a 2+ to 5+ charge state. The total cycle time was fixed to 3.3 s. The Q2 transmission window was 100 Da for 100 %. Four time bins were summed for each scan at a pulse frequency value of 11 kHz by monitoring the 40-GHz multichannel time-to-digital converter detector with a four-anode channel detector ion. A sweeping collision energy setting of 35 ± 5 eV coupled with iTRAQ adjust rolling collision energy was applied to all precursor ions for collision-induced dissociation. Dynamic exclusion was set for half the peak width (15 s), and then the precursor was refreshed off the exclusion list.

#### Proteomics data analysis

Raw data files were converted into MGF files by the Proteome Discoverer 1.2 (Thermo, USA), and the MGF files were searched. Proteins were identified by using a Mascot search engine (Version 2.3.02, Matrix Science, UK). The parameters were set as follows: peptide tolerance, 0.05 Da; fragment MS tolerance, 0.1 Da; fixed modification, methylthio (C); and variable modifications involving acetyl (N-term), pyro-glu (N-term Q), oxidation (M), deamidation (N, Q), and iTRAQ 8-plex (N-term, K, Y). A maximum of one missed cleavage was allowed in the trypsin digests, and peptide charge states were set to +2 and +3. The search that was performed in Mascot was an automatic decoy database search. To reduce the probability of false positives, raw spectra from the actual database were tested against a generated database of random sequences. Only peptides with scores (> 20) significant at the 95 % confidence level were considered to be reliable and were used for protein identification. Each identified protein involves at least one unique peptide. For protein quantitation, a protein was required to contain at least two unique peptides. Protein quantitative ratios were weighted and normalized relative to the median ratio in Mascot. Only the proteins with significant quantitative ratios between the two treatments with fold changes >1.2 and *P*-values < 0.05 were considered to be differentially expressed and significant.

### Integrated analysis of the immunomodulatory effects of *H. perezi*

To further explore the effects of *H. perezi* on the crustacean immune response, the immune-related pathways influenced by the parasite were systematically analyzed via the KEGG enrichment analysis of the differentially expressed miRNAs (DEMs) and proteins (DEPs) involved in the immune responses to the parasite. The concordance of the immune related KEGG pathways enriched from the two omics data was evaluated. The putative target genes of the DEMs were searched against the identified proteins from the proteomics data. The pairs of the DEMs-target proteins were selected to further investigate the integrative immune-regulatory effects induced by *Hematodinium* infection.

### Validation of Representative DEMs and DEPs with qRT-PCR Assays

To verify the reliability of the sequencing data, the transcriptional profiles of 10 DEMs and 10 DEPs involved in critical host immune pathways (ex. Phagosome, Complement and coagulation cascades, Focal adhesion, Peroxisome, Lysosome, and ECM-receptor interaction) and displaying significant alterations were further analyzed by qRT-PCR. Total RNAs were isolated from the same subset of hemocytes samples collected at the different time points using Trizol reagent (TaKaRa, Japan). The extracted RNAs were treated with RNase-free DNase I (TaKaRa, Japan) to remove contaminating genomic DNAs. The transcripts of the DEMs and DEPs were analyzed and validated by qRT-PCR assays using the Mir-X miRNA qRT-PCR SYBR^®^ Kit (TaKaRa, Japan) and the PrimeScript^TM^ RT reagent Kit (TaKaRa, Japan) as described in previous studies (Li et al., 2018, 2019) [45, 46]. The primers used for the qRT-PCR assays are listed in S1 & S2 Tables. The reverse primer (mRQ 3′ primer) and the primers of the U6 snRNA used in the qRT-PCR assays were provided in the Mir-X miRNA qRT-PCR SYBR® Kit (Takara, Japan). The qRT-PCR data were analyzed by the 2^−ΔΔCt^ method as described in Livak & Schmittgen (2001) [106]. Data were shown as the mean ± standard deviation (SD) and analyzed via the one-way analysis of variance (ANOVA), followed by Duncan’s multiple range test using the SPSS Statistics software (version 20.0, SPSS, USA). Differences were considered significant when *P* < 0.05.

## Author Contributions

ML and CL conceived and designed the experiments, ML, QH and XL performed the experiments and analyzed the data, HJS advised on data analysis, ML wrote the original draft, HJS and CL revised the manuscript. All authors reviewed and approved the final manuscript.

## Acknowledgements

The hospitality and assistance of Mr. Fei Wang and his colleagues at Qingdao Yi Hai Feng Aquatic Products Co., Ltd are gratefully appreciated. We are particularly grateful to the other team members in the laboratory and the team of Prof. Zhiming Yu at IOCAS for their laboratory assistance. We also thank the anonymous reviewers for their constructive criticism and suggestions on the manuscript. This paper is Contribution Number XXXX of the Virginia Institute of Marine Science, William & Mary.

## Supporting information

**S1 Fig.** Length distribution of the identified unique miRNAs.

**S2 Fig.** Venn diagram of the identified miRNAs. MiRNAs were identified in hemocytes of *Portunus trituberculatus* at different time-points after challenge with *Hematodinium perezi* (4, 8 and 16 days), and compared to miRNAs from hemocytes from uninfected crabs (0 day). The numbers of the identified unique miRNAs between the time points are indicated.

**S3 Fig.** Summary of the proteome data in hemocytes of *P. trituberculatus* by iTRAQ. Total spectra are the secondary mass spectrums, and spectra are the secondary mass spectrums after quality control. Unique Peptide is the identified peptides which belong only to a group of proteins, and protein is identified by Mascot 2.3.02 software.

---

**S1 Table.**
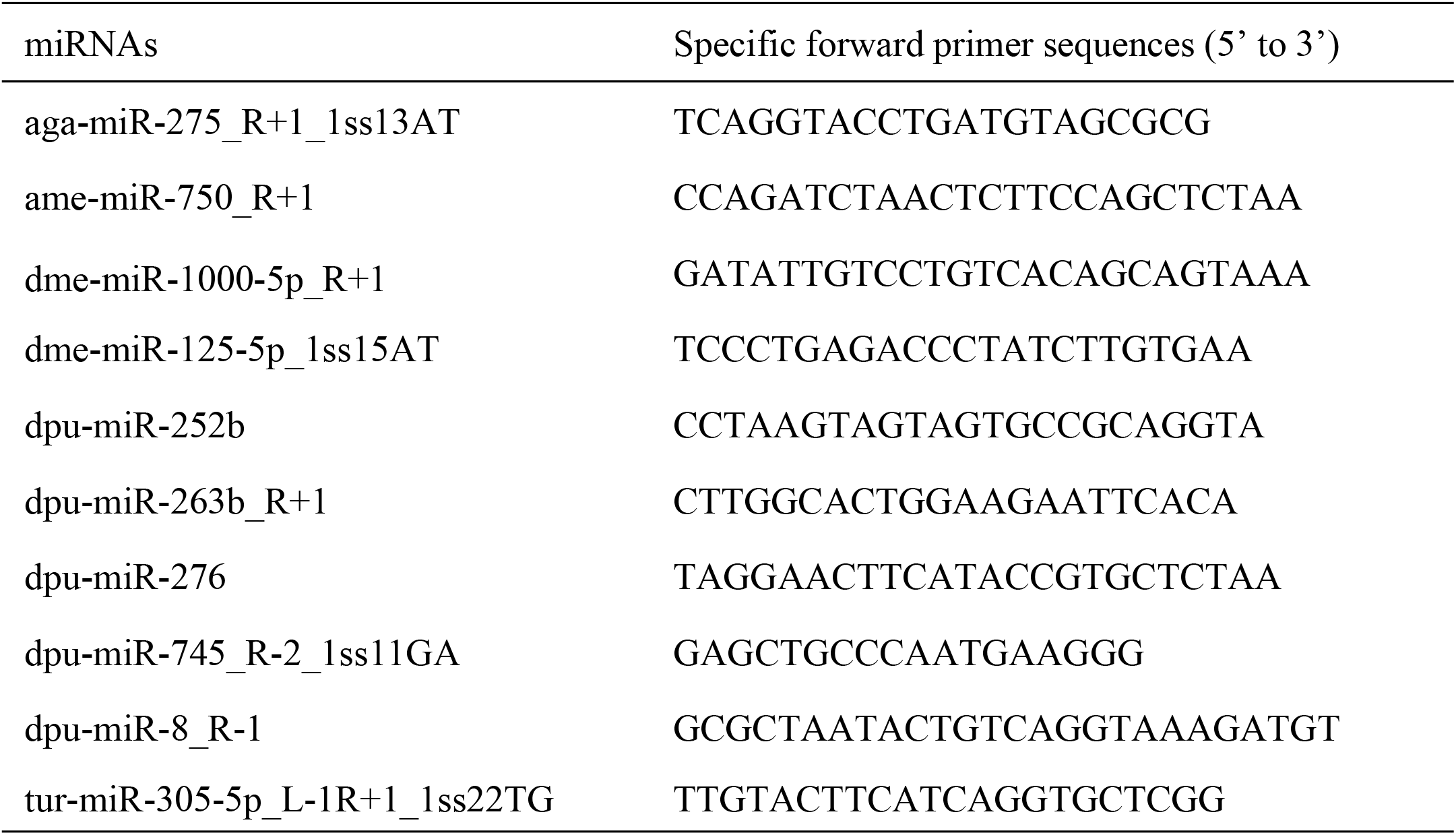
Primers designed for qRT-PCR assessment of select miRNAs expression.

**S2 Table.**
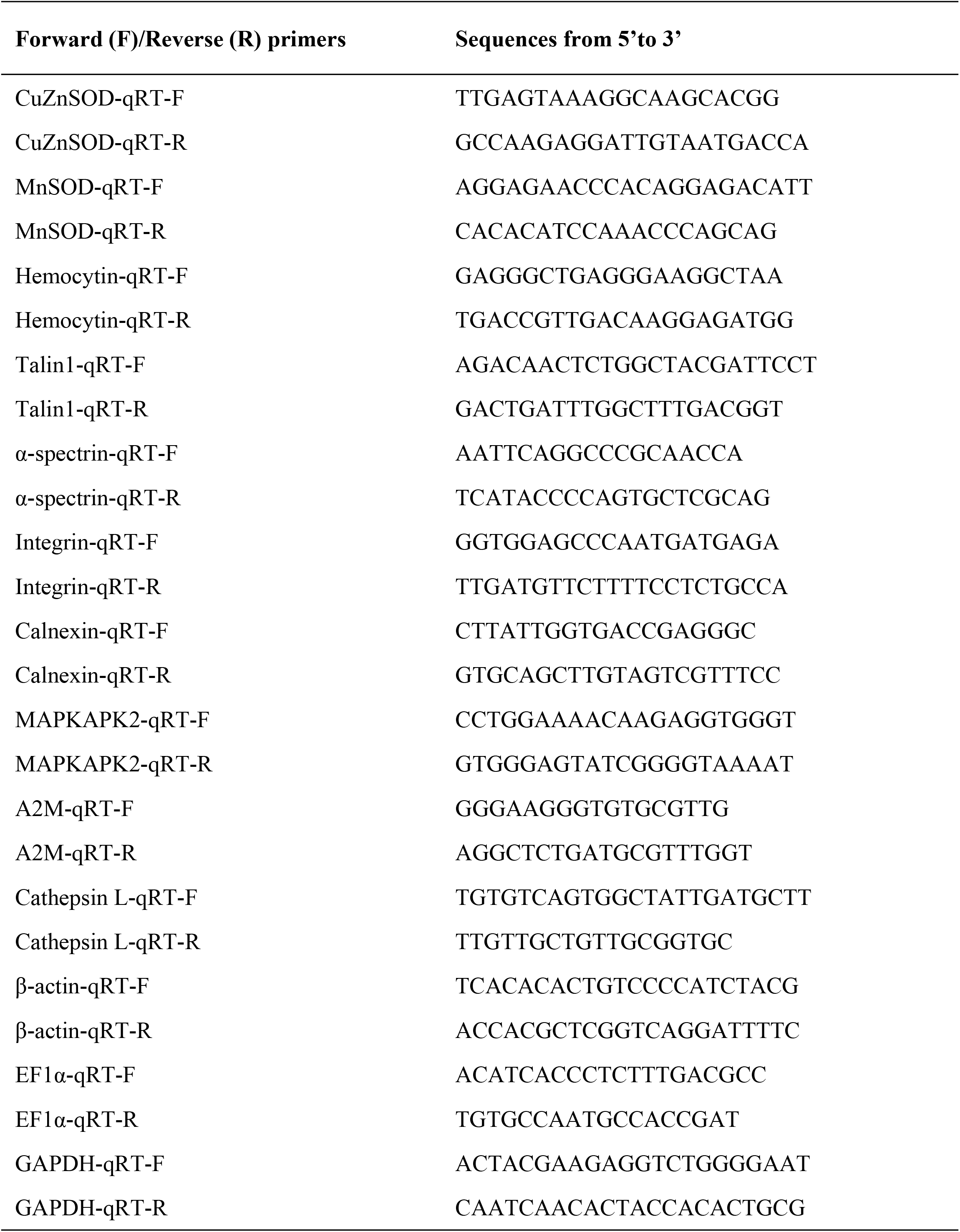
Primers designed for qRT-PCR assessment of select proteomic transcripts of DEPs.

